# Investigating Population-scale Allele Specific Expression in Wild Populations of *Oithona similis* (Cyclopoida, Claus 1866)

**DOI:** 10.1101/599076

**Authors:** Romuald Laso-Jadart, Kevin Sugier, Emmanuelle Petit, Karine Labadie, Pierre Peterlongo, Christophe Ambroise, Patrick Wincker, Jean-Louis Jamet, Mohammed-Amin Madoui

## Abstract

Allele-specific expression (ASE) is a widely studied molecular mechanism at cell, tissue and organism levels. Here, we extrapolated the concept of ASE to the population-scale (psASE), aggregating ASEs detected at smaller scales. We developed a novel approach to detect psASE based on metagenomic and metatranscriptomic data of environmental samples containing communities of organisms. This approach which measures the deviation between the frequency and the relative expression of biallelic loci, was applied on samples collected during the *Tara* Oceans expedition (2009-2013), in combination to new *Oithona similis* transcriptomes, a widespread marine copepod. Among a total of 25,768 single nucleotide variants (SNVs) of *O. similis*, 587 (2.3%) were targeted by psASE in at least one population. The distribution of SNVs targeted by psASE in different populations is significantly shaped by population genomic differentiation (p-value = 9.3×10^−9^), supporting a partial genetic control of psASE. To investigate the link between evolution and psASE, loci under selection were compared to loci under psASE. A significant amount of SNVs (0.6%) were targeted by both selection and psASE (p-values < 9.89×10^−3^), supporting the hypothesis that natural selection and ASE may lead to the same phenotype. Population-scale ASE offers new insights into the gene regulation control in populations and its link with natural selection.

## Introduction

Allele-specific expression (ASE), or allelic imbalance, refers to the difference of expression between two alleles of a locus in a heterozygous genotype. Recently, several studies led to a better understanding of ASE thanks to the development of advanced tools (Skelly et al. 2011; Mayba et al. 2014; Castel et al. 2015; Harvey et al. 2015; Lu et al. 2015; Rivas et al. 2015; Miao et al. 2018) allowing its detection at cellular (Milani et al. 2009; Ginart et al. 2016; Huang et al. 2017; Dong & Jiang 2019) tissue (McKean et al. 2016; Zhuo et al. 2017; Liu et al. 2018) and/or inter-individual levels (Josephs et al. 2015; Tung et al. 2015; Lonsdale et al. 2017; Wang et al. 2017). Moreover, studies began to question the relative contribution of genetics and environment on gene expression using ASE in human (Cheung et al. 2008; Buil et al. 2014; Cheung et al. 2008; Moyerbrailean et al. 2016; Knowles et al. 2017) and fruit flies (Leon-Novelo et al. 2017). Because the availability of reference genomes, numerous individual RNA-seq and whole-genome genotyping data still constitutes an important challenge for many species, there are few studies investigated ASE in natural populations (Tung et al. 2015; Wang et al. 2017).

ASEs are mainly due to epigenetic or genetic processes in *cis*-regulatory variations. Through DNA methylation, chromatin state or histone modifications, epigenetics could repress a disadvantageous or a specific parental allele, leading in some cases to monoallelic expression, as demonstrated in a variety of organisms including *Arabidopsis thaliana*, mouse, maize or bumblebee (Szabo & Mann 1995; Zhang & Borevitz 2009; Wei & Wang 2013; Ginart et al. 2016; Lonsdale et al. 2017). ASE may also have a genetic origin through, for example, mutations in transcription factor binding sites (Bailey et al. 2015; Cavalli et al. 2016), or post-transcriptional mechanisms like non-sense mediated decay (Castel et al. 2015; Rivas et al. 2015; Pirinen et al. 2015).

However, ASE at population-scale (psASE) has never been investigated. At this level, psASE may aggregate classical ASE at cell, tissue or individual levels (Figure 1a), but also differential expression between two homozygous genotypes. However, measurement of psASE would require the sequencing of several individuals at genomic and transcriptomic levels separately. To measure psASE on small organisms, an alternative approach could be to take advantage of metagenomic and metatranscriptomic data that already contains sequences from pooled individuals. In this context, intra-species variants of a single species have to be extracted, allowing to evaluate if the population-scale relative expression of an allele deviates from its genomic frequency (Figure 1b).

**Figure 1:**
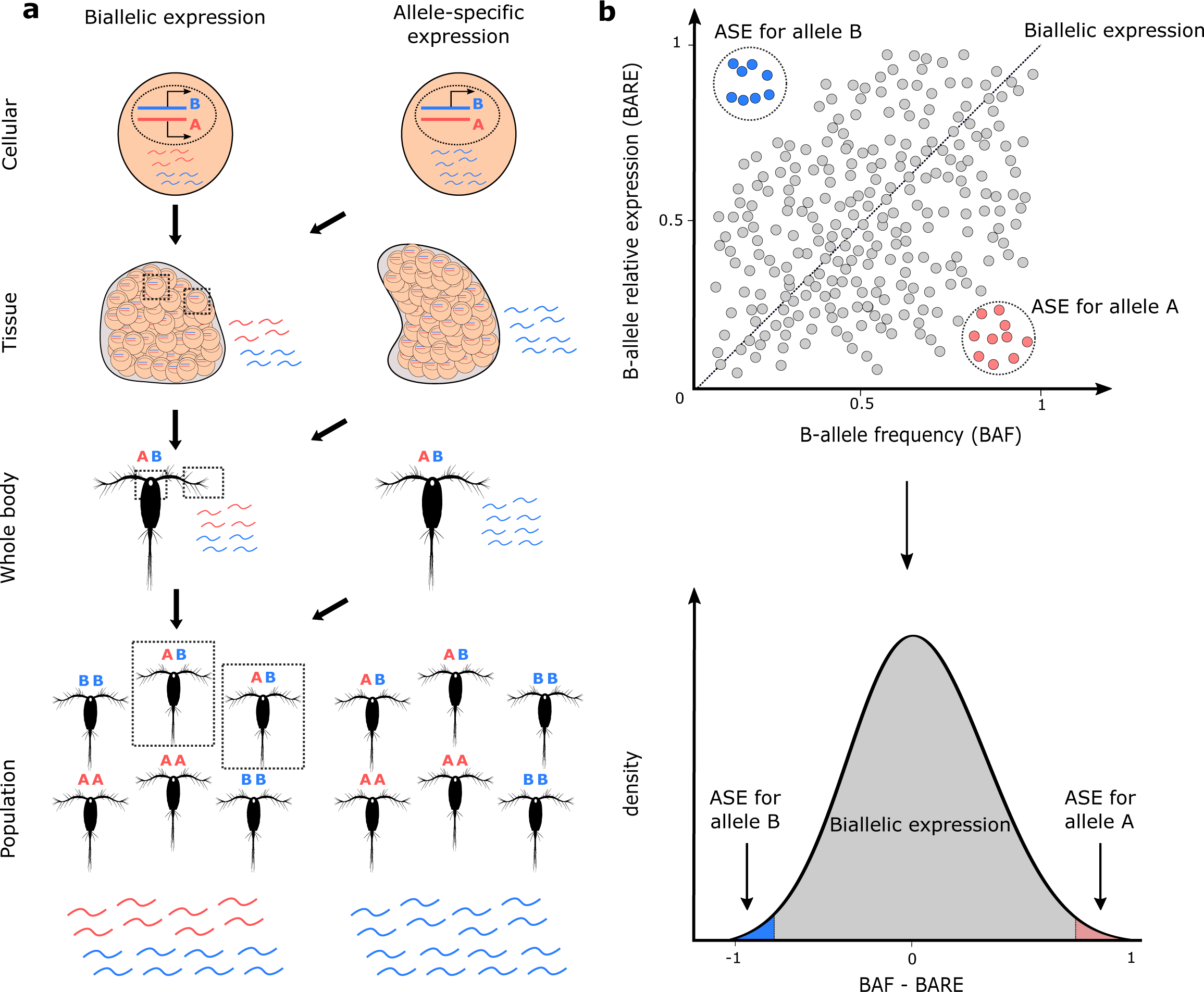
Allele-specific expression at population-scale. **a**, Different scales of allele-specific expression detection for a heterozygous gene, from population to cellular levels. For a heterozygous genotype, ASE is understood as the difference in expression between to alleles of a single gene, opposed to strict biallelic expression. For clarity, the example of ASE presented here is monoallelic expression. **b**, Null hypotheses to detect psASE. On the upper panel, BAF and BARE are plotted for each variant (points). In grey, the majority of variants, with allele relative expression following genomic frequency. In red (blue), variants under potential psASE for allele A (B). The black dotted line stands for the theoretical regression curve reflecting proportionality between BAF and BARE. On the lower panel, the deviation between BAF and BARE is computed and should theoretically follow Gaussian distribution centered on 0. In the grey areas, variants with no apparent deviation between BAF and BARE; in red (blue) area, variants in potential psASE for allele A (B), with a significant difference between BAF and BARE.

Copepods, and particularly species belonging to the *Oithona* genus, are small crustaceans forming the most abundant metazoan on Earth (Humes 1994; Gallienne 2001; Kiørboe 2011). This abundance, reflecting strong adaptive capacities to environmental fluctuations, together with large hypothetic effective population size (Peijnenburg & Goetze 2013; Riginos et al. 2016; Madoui et al. 2017; Arif et al. 2018) make this species suitable for population genomics analyses. In addition, they play an ecological key role in biogeochemical cycles and in the marine trophic food chain (Wassmann et al. 2006). In this study, we propose to test our model to detect psASE by focusing on the widespread epipelagic copepod, *Oithona similis* (Cyclopoida, Claus 1866). We used environmental samples collected by the *Tara* Oceans expedition (Karsenti et al. 2011; Pesant et al. 2015) during its Arctic phase (2013), an area where *O. similis* is known to be highly abundant (Blachowiak-Samolyk et al. 2008; Dvoretsky 2012; Zamora-Terol et al. 2013; Castellani 2016) and for which both metagenomic and metatranscriptomic data are available. Variants of *O. similis* were extracted and tested for psASE. In second time, we tried to decipher the potential link between psASE, genomic differentiation and natural selection.

## Material and Methods

### Variant calling using *Tara* Oceans metagenomic and metatranscriptomic data

We used metagenomic and metatranscriptomic reads generated from samples of the size fraction 20–180 µm collected in seven *Tara* Oceans stations (TARA_155, 158, 178, 206, 208, 209 and 210)(Figure 2a) according to protocols described by Alberti et *al.* (Supplementary Table S2). Because of the lack of a reference genome, the reference-free variant caller *DiscoSNP++* (Uricaru et al. 2014; Peterlongo et al. 2017) was used to extract SNVs simultaneously from raw metagenomic and metatranscriptomic reads, and was ran using parameter –b 1. Only SNVs corresponding to biallelic loci with a minimum of 4x of depth of coverage in all stations were initially selected. Then, to capture loci belonging to *Oithona similis*, SNVs were clustered based on their loci co-abundance across samples using density-based clustering algorithm implemented in the R package dbscan (Ester et al. 1996; Ram et al. 2010) and ran with parameters epsilon = 10 and minPts = 10. This generated three SNVs clusters, the largest of which contained 102,258 SNVs. To ensure only the presence of *O. similis* SNVs, we observed the fitting of the depth of coverage to the expected negative binomial distribution in each population (Supplementary Figure S2). As the variant calling step is reference-free, the alternative allele (here, B-allele) is arbitrary set by *DiscoSNP++*. For each variant in each population, the B-allele frequency (BAF) and the population-level B-allele relative expression (BARE) were computed as follow, 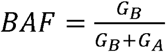 and 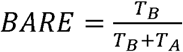, with *G*_*A*_, *G*_*B*_ the metagenomic read counts supporting the reference and alternative alleles respectively and *T*_*A*_, *T*_*B*_ the metatranscriptomic read counts supporting the reference and alternative alleles respectively.

**Figure 2:**
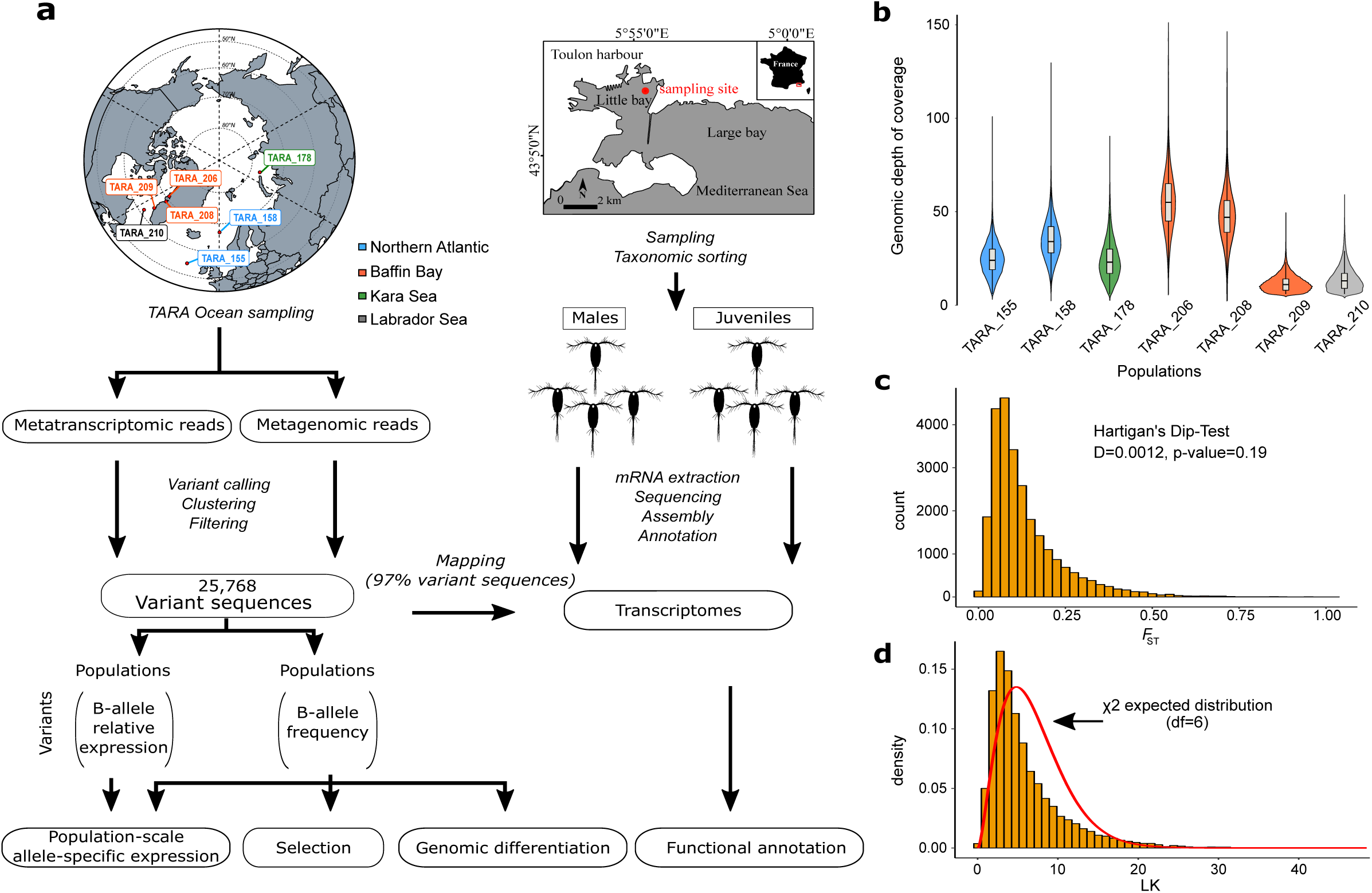
Genomic variation data of *O. similis.* **a**, Scheme representing the study, from samplings to analyses **b**, Genomic depth of coverage distributions of the set of 25,768 variants by sample **c**, *F*_ST_ distribution across the seven samples **d**, LK distribution. The red line represents χ^2^ theoretical distribution (df = 6).

Biallelic loci were then filtered based on their metagenomic coverage. For each sample, the median and standard deviation σ of the distribution of metagenomic coverage of all biallelic loci were estimated. Biallelic loci must be characterized by a metagenomic coverage between a limit of median ± 2σ, with a minimum and maximum of 5x and 150x coverage in each sample to avoid low covered and multicopy genomic regions. To keep out rare alleles and potential calling errors, only variants characterized by a BAF comprised between 0.9 and 0.1, and a BARE between 0.95 and 0.05 in at least one population were chosen for the final dataset resulting in 25,768 biallelic loci.

To ensure that these loci belong to *O. similis*, the global *F*-statistics (or Wright’s fixation index (Wright 1951; B. S. Weir and C. Clark Cockerham 1984)) over the seven populations was computed as follow, 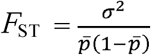, with 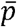 being the mean allele frequency across all the samples, and its distribution was tested for unimodality via a Hartigans’ dip test (Hartigan & Hartigan 1985). Moreover, LK statistics (R.C. & J. 1973) was computed as follow, 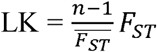, and compared to the expected chi-squared distribution with df = n-1, with n being the number of populations and 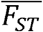 being the mean *F*_*ST*_ across all loci.

### Population-scale ASE detection using metagenomic and metatranscriptomic data

In each population, we first selected variants for BAF ≠ {0,1}. We tested the correlation between BAF and BARE and then we computed *D = BAF-BARE*, as the deviation between the BAF and the BARE. In the seven samples, *D* followed a Gaussian distribution centered on 0, allowing the estimation of the Gaussian distribution parameters and test the probability of a variant to belong to this distribution (this is later referred here as “*D*-test” or “deviation test”). Given the large number of tests, we applied the Benjamini and Hochberg approach (Benjamini & Hochberg 1995) to control the False Discovery Rate (FDR). We also computed a “low expression bias” test by comparing the read counts *T*_*A*_ and *T*_*B*_ to the observed metagenomic proportion 1-BAF and BAF respectively with a chi-squared test and applied the Benjamini and Hochberg correction for multiple testing.

The deviation and the low expression bias tests were applied to all loci in the seven populations separately. Loci with a deviation test q-value < 0.1 and a low expression bias test q-value < 0.1 were considered under psASE. This generates seven sets of candidate loci under psASE, one set for each population.

### Estimation of genomic differentiation and detection of variants under selection

Pairwise-*F*_ST_ was computed for each locus between each pair of population and the median pairwise-*F*_ST_ was retained to measure the genomic differentiation between each population. A Mantel test was performed to test for isolation-by-distance between median pairwise-*F*_ST_ and geographic Euclidean distances using *vegan* v2.5-2 (Oksanen et al. 2018) and *geosphere* v1.5-7 (Hijmans 2017) R packages. The *pcadapt* R package v4.0.2 (Luu et al. 2017) was used to detect selection among populations from the B-allele frequency matrix. The computation was run on “*Pool-seq*” mode, with a minimum allele frequency of 0.05 across the populations, and variants with a corrected Benjamini and Hochberg p-value < 0.05 were considered under selection.

### Modelling psASE with genomic differentiation

The seven sets of candidate loci under psASE were crossed to identify variants under psASE in several populations (named “shared psASEs”) and all non-empty crossings between populations were represented by an upset plot.

To test if populations characterized by weak genetic differentiation tend to share more loci under psASE than genetically distant populations, we modelled the number of shared psASEs between populations by genomic differentiation using a non-linear model: *E(Y)* ∼ *a exp(bX) + c*, with Y being the number of shared ASE and X the genomic differentiation. The latter was estimated by computing the median-*F*_ST_ of each non-empty crossing. To find the starting values, the model was linearized as follow, *log(E(Y)*−*c0)* ≈ *log(a)+bX*, with *c0 = min(Y) * 0.5* and *a* and *b* parameters were estimated via the *lm* R function. The non-linear model was then applied, and least squares estimates were used via the *nls* R function. Pearson correlation between the fitted and empirical values was then computed via the *cor.test* R function.

### psASE and link with natural selection

To identify alleles targeted by both psASE and natural selection, the set of variants under ASE in each population was crossed with the set of loci detected under selection. The size of the intersection was tested by a hypergeometric test, Hypergeometric(q,m,n,k), with *q* being number of alleles under ASE in the population and under selection (size of intersection), *m* being the total number of alleles under selection, *n* being the total number of variants under neutral evolution, and *k* being the total number of alleles under psASE in the tested population. We considered that, in a given population, the number of alleles under both psASE and selection was significantly higher than expected by chance for p-value < 0.05.

### Material sampling, mRNA extraction and Mediterranean *O. similis* transcriptomes sequencing

To conduct a functional analysis, Mediterranean *O. similis* transcriptomes were produced. *Oithona similis* specimens were sampled at the North of the Large Bay of Toulon, France (Lat 43°06’ 02.3” N and Long 05°56’ 53.4” E). Sampling took place in November 2016. The samples were collected from the upper water layers (0-10m) using zooplankton nets with a mesh of 90µm and 200µm (0.5 m diameter and 2.5 m length). Samples were preserved in 70% ethanol and stored at -4°C. From the Large Bay of Toulon samples, *O. similis* individuals were isolated under the stereomicroscope (Rose 1933; Nishida 1985). We selected two different development stages: four copepodites (juveniles) and four adult males. Each individual was transferred separately and crushed, with a tissue grinder (Axygen) into a 1.5 mL tube (Eppendorf). Total mRNAs were extracted using the ‘RNA isolation’ protocol from NucleoSpin RNA XS kit (Macherey-Nagel) and quantified on a Qubit 2.0 with a RNA HS Assay kit (Invitrogen) and on a Bioanalyzer 2100 with a RNA 6000 Pico Assay kit (Agilent). cDNA were constructed using the SMARTer-Seq v4 Ultra low Input RNA kit (ClonTech). The libraries were built using the NEBNext Ultra II kit, and were sequenced with an Illumina.

### Transcriptomes assembly and annotation

Each read set was assembled with Trinity v2.5.1 (Haas et al. 2013) using default parameters and transcripts were clustered using cd-hit v4.6.1 (Fu et al. 2012) (Supplementary Table S1). To ensure the classification of the sampled individuals, each ribosomal read set were detected with SortMeRNA (Kopylova et al. 2012) and mapped with bwa v0.7.15 using default parameters (Li & Durbin 2009) to 82 ribosomal 28S sequences of *Oithona* species used in Cornils et al., 2017 (Supplementary Figure S1). The transcriptome assemblies were annotated with Transdecoder v5.1.0 (Haas et al. 2013) to predict the open reading frames (ORFs) and protein sequences (Supplementary Table S1). In parallel, homology searches were also included as ORF retention criteria; the peptide sequences of the longest ORFs were aligned on *Oithona nana* proteome (Madoui et al. 2017) using DIAMOND v0.9.22 (Buchfink et al. 2014). Protein domain annotation was performed on the final ORF predictions with Interproscan v5.17.56.0 (Jones et al. 2014) and a threshold of e-value <10^−5^ was applied for Pfam annotations. Finally, homology searches of the predicted proteins were done against the nr NCBI database, restricted to Arthropoda (taxid: 6656), with DIAMOND v0.9.22.

### Variant annotation

The variant annotation was conducted in two steps. First, the variant sequences were relocated on the previously annotated *O. similis* transcripts using the “VCF_creator.sh” program of *DiscoSNP++*. Secondly, a variant annotation was carried out with SNPeff (Cingolani et al. 2012) to identify the location of variants within transcripts (i.e., exon or UTR) and to estimate their effect on the proteins (missense, synonymous or nonsense). The excess of candidate variant annotations was tested in the following classes: missenses, synonymous, 5’ and 3’UTR. A significant excess was considered for a hypergeometric test p-value < 0.05.

### Gene enrichment analysis

To identify specific biological function or processes associated to the variants, a domain-based analysis was conducted. The Pfam annotation of the transcripts carrying variants targeted by ASE and selection was used as entry for dcGO Enrichment (Fang & Gough 2013). A maximum of the best 300 GO-terms were chosen based on their z-score and FDR p-value (<10^−3^) in each ontology category. To reduce redundancy, these selected GO-terms were processed using REVIGO (Supek et al. 2011), with a similarity parameter of 0.5 against the whole Uniprot catalogue under the SimRel algorithm. To complete the domain-based analysis, the functional annotations obtained from the homology searches against the nr were manually curated.

## Results

### Extracting polar *Oithona similis* variants from environmental samples

From metagenomic and metatranscriptomic raw data of seven sampling stations (Figure 2a), we identified 102,258 variants using a reference-free approach. Among them, 25,768 expressed *O. similis* variants were retrieved after filtering. To ensure that the variants belonged to *O. similis*, we performed three different analyses. First, in each sample, the distributions of variable loci depth of coverage were unimodal (Figure 2b) and fitted the expected negative binomial distributions (Supplementary Figure S2). Second, 97% of 25,768 variants were relocated on Mediterranean *O. similis* transcriptomes (Figure 2a). Third, the global distribution of *F*_ST_ of the seven populations was unimodal (Hartigans’ dip test, D=0.0012, p-value=0.19) with a low median *F*_ST_ at 0.1 (Figure 2c), confirmed by the pairwise-*F*_ST_ distributions (Supplementary Figure S3d). Finally, the LK distribution over all the loci followed the expected chi-squared distribution (Figure 2d), showing that most of the loci follow the neutral evolution model, as expected in a single species.

### *Oithona similis* genomic differentiation in Arctic Seas

The seven populations were globally characterized by a weak to moderate differentiation, with a maximum median pairwise-*F*_ST_ of 0.12 between populations from TARA_210 and 155/178 (Figure 3b, Supplementary Figure S3). Populations from stations TARA_158 (Norway Current), 206 and 208 (Baffin Bay) were genetically closely related, with the lowest median pairwise-*F*_ST_ (0.02), despite TARA_158 did not co-geolocalize with the two other stations. The four other populations (TARA_155, 178, 209, and 210) were equally distant from each other (0.1-0.12). Finally, TARA_158, 206 and 208 on one side, and TARA_155, 178, 210 and 209 on the other side showed the same pattern of differentiation (0.05-0.07). A Mantel test was performed and revealed no correlation between *F*_ST_ and geographic distances (r = 0.34, p-value = 0.13) (Supplementary Figure S5).

**Figure 3:**
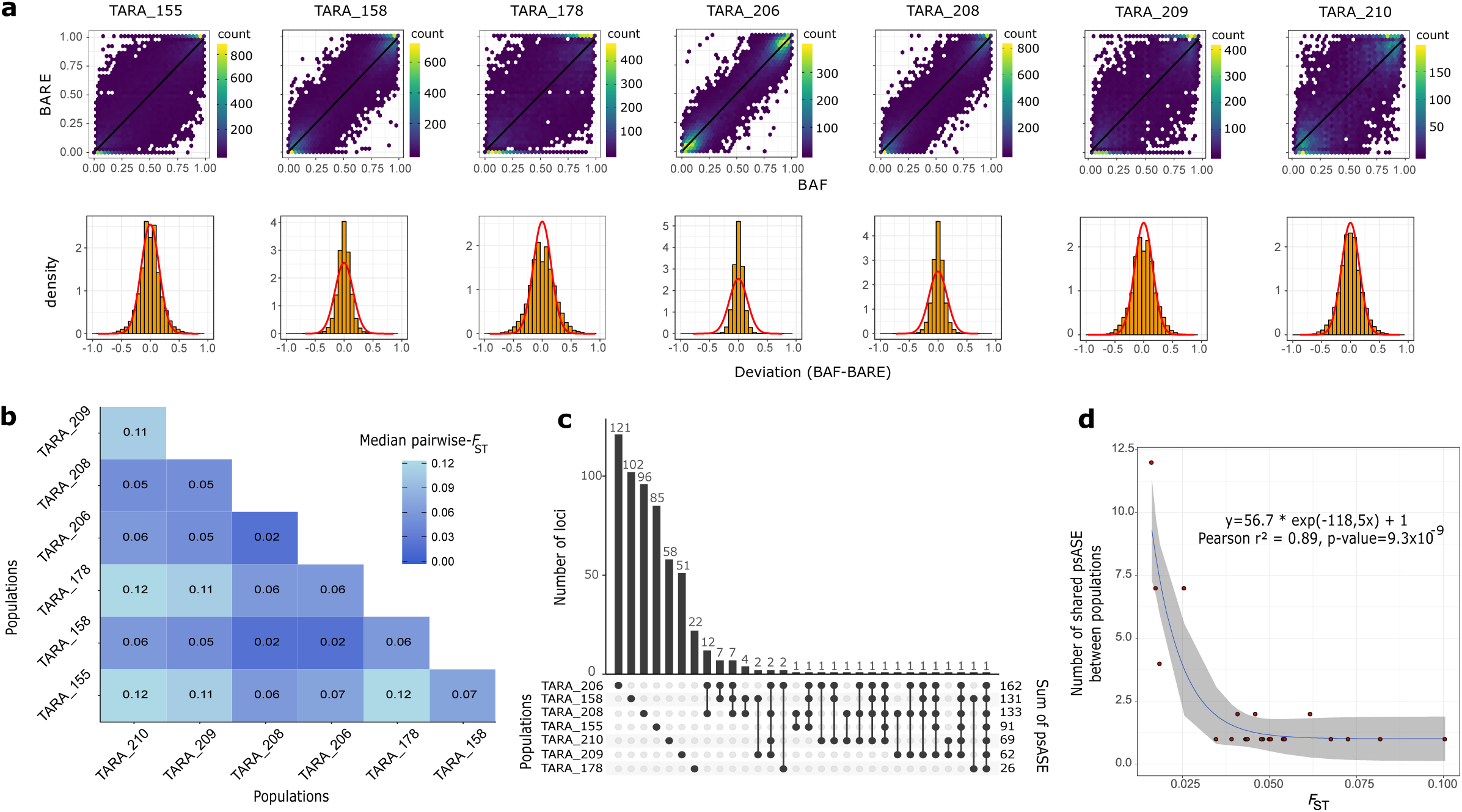
Population-scale allele-specific expression and its link with genomic differentiation. **a**, Each column corresponds to a population. Upper panel represents the relation between BAF and BARE, each hexagone corresponds to an area containing the number of variants indicated by the color scale. Black lines are the linear regression curves. Lower panel represents the deviation *D* distribution. The red lines correspond to the Gaussian distribution estimated from the data. **b**, Pairwise-*F*_ST_ matrix. The median (mean) of each pairwise-*F*_ST_ distribution computed on allele frequencies is indicated **c**, Upset plot of psASE detection in the seven populations. Each bar of the upper plot corresponds to the number of variants under psASE in the combination of population(s) indicated by black dots in the lower plot. **d**, Genomic differentiation and shared psASE. Each dot is a combination of population as presented in the lower panel of the upset plot. The blue line represents the non-linear regression curve estimated from the data and 95% confidence interval in grey.

### Detection of population-scale ASE

The number of SNVs tested for psASE varied between 13,454 and 22,578 for TARA_210 and 206 respectively. As expected, most of the loci presented a strong correlation between B-allele frequency and B-allele relative expression (Figure 3a, Table 1). Thus, we measured the *D* deviation, representing the deviation between BAF and BARE, which followed a Gaussian distribution in each population (Figure 3a). Variants under psASE (i.e. having a *D* significantly higher or lower than expected and passing the low expression bias test) were found in every population, ranging from 26 to 162 variants for TARA_178 and 206 respectively (Table 1). Overall, we found 587 variants under psASE, including 535 population-specific ASEs, and 52 ASEs shared by several populations (Figure 3c). Remarkably, 30 ASEs out of the 52 were present in the populations from TARA_158, 206 and 208 that correspond to the genetically closest populations. By comparing the number of shared psASEs in the different sets of populations to their genomic differentiation, we found a negative trend between the two (with a strong negative *b* estimate), illustrated by a significant correlation between non-linear fitted and empiric values (0.89, p-value 9.3×10^−9^, Figure 3d). This modelling shows that genetically close populations tend to share the same variants under psASE.

**Table 1:** Allele-specific expression detection and link with selection by population. Adjusted-r^2^ corresponds to linear regression performed between BAF and BARE of all tested loci in the respective populations.

| Population | Genomic median<br>depth of coverage | Number of tested<br>variants | Adjusted-r<sup>2</sup> | Number of<br>variants under<br>ASE | Number of variants<br>under ASE and selection | Hypergeometric<br>test p-value |
| --- | --- | --- | --- | --- | --- | --- |
| TARA_155 | 25 | 18,812 | 0.77 | 91 | 6 | 9.89x10<sup>-3</sup>* |
| TARA_158 | 35 | 21,476 | 0.91 | 131 | 29 | 5.06x10<sup>-20</sup>* |
| TARA_178 | 24 | 18,145 | 0.67 | 26 | 9 | 5x10<sup>-11</sup>* |
| TARA_206 | 55 | 22,578 | 0.95 | 162 | 42 | 4.82x10<sup>-31</sup>* |
| TARA_208 | 48 | 21,469 | 0.93 | 133 | 14 | 2.2x10<sup>-6</sup>* |
| TARA_209 | 12 | 13,956 | 0.73 | 62 | 24 | 1.05x10<sup>-23</sup>* |
| TARA_210 | 14 | 13,454 | 0.71 | 69 | 42 | 8.89x10<sup>-51</sup>* |
| Overall | - | 25,768 | - | 587 (2.3%) | 152 (0.59%) | - |

### Loci targeted by population-level ASE and selection in Arctic populations

The set of variants was tested for selection using *pcadapt*. The PCA decomposed the genomic variability in six components (Supplementary Figure S3a,b,c); the first two components discriminated TARA_155 and 178 from the others (32% and 28.1% variance explained respectively), and the third component differentiated TARA_210 and 209 (19.5%). The fourth principal component separated TARA_209 and 210 from 158/206/208 (11.3 %), with the last two concerning TARA_158/206/208. Globally, these results dovetailed with the *F*_ST_ analysis, with details discussed later. Finally, we detected 674 variants under selection, representing 2.6% of the dataset (corrected p-value < 0.05).

The seven sets of variants under psASE were crossed with the set of variants under selection (Figure 4a). The size of the intersections ranged from 6 to 42 variants (TARA_155 and 210/206) and was significantly higher than expected by chance for all the populations (Figure 4b, hypergeometric test p-value < 0.05). It represented a total of 152 unique variants under selection and ASE in at least one population, corresponding to 23% and 26% of variants under psASE and under selection respectively.

**Figure 4:**
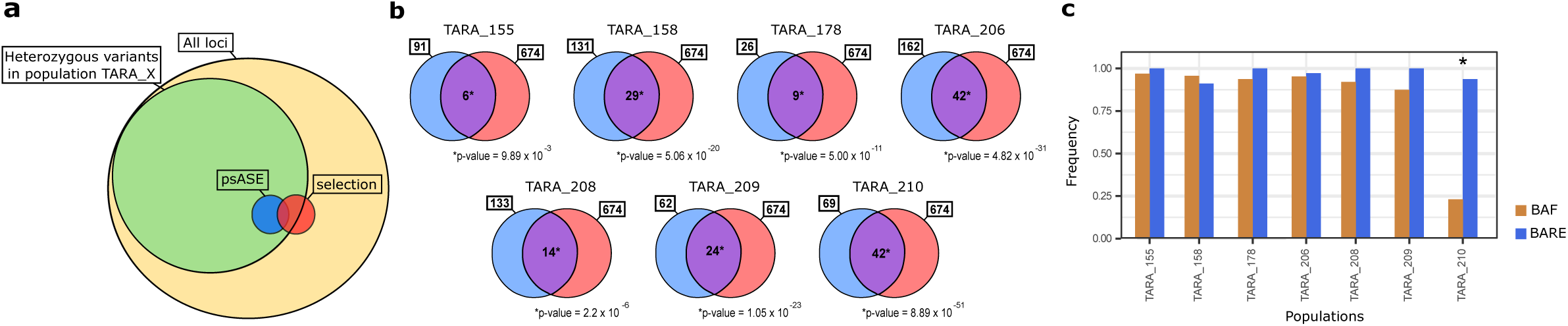
Crossing psASE and selection. **a**, Scheme representing the method; in yellow the total dataset of SNVs, in green the tested SNVs in one population, in blue SNVs under psASE in this population, in red the SNVs targeted by selection. **b**, Crossing the candidate variants under psASE (blue) and those under selection (red) for each population. The hypergeometric p-value corresponds to the significance of the amount of variants both under psASE in the considered population and under selection (purple). **c**, Metagenomic and metatranscriptomic profiles of the SNV 1219525, under psASE in TARA_210 (*) and reaching near fixation in the other populations. BAF and BARE for each population in golden and blue respectively.

### Functional analysis of genes targeted by population-scale ASE and selection

Among the 152 loci targeted by psASE and selection, 145 were relocated on *O. similis* transcripts (Supplementary Table S4). Amid these transcripts, 137 (90%) had a predicted ORF, 97 (64%) were linked to at least one Pfam domain and 90 (59%) to a functional annotation. Fifteen SNVs were missense variations, 59 synonymous, 31 and 29 were in 5’ and 3’ UTR, without any significant excess (Supplementary Table S4 and S5). Based on homology searches (Supplementary Table S4), eight genes were linked to nervous system (Table 2). Among them, two genes were involved in glutamate metabolism (omega-amidase NIT2 and 5-oxoprolinase), three were predicted to be glycine, γ-amino-butyric acid (GABA) and histamine neuroreceptors. Finally, four were also implicated in arthropods photoreceptors. The domain-based analysis confirmed these results, with an enrichment in GO-terms biological process also linked to nervous system (Supplementary Figure S6).

**Table 2:** Functional annotations of variants targeted by ASE and selection implicated in nervous system.

| VarID | Ref | Alt | Homology search | Pfam | SnpEff<br>Localization | SnpEff<br>Impact | References |
| --- | --- | --- | --- | --- | --- | --- | --- |
| 722267 | A | G | histamine H1 receptor | PF00001 | 3 ' UTR | MODIFIER | (Monastirioti 1999;<br>Stuart 1999; Denno et<br>al. 2016) |
| 9665345 | T | G | chaoptin | PF13306 \| PF13855 | synonymous<br>variant | LOW | (Krantz & Zipursky<br>1990; Gurudev et al.<br>2014) |
| 15623788 | G | A | eye-specific<br>diacylglycerol kinase | PF13637 | synonymous<br>variant | LOW | (Masai et al. 1993;<br>Wang & Montell 2007) |
| 23795359 | A | T | vang-like protein 2-B | PF06638 | synonymous<br>variant | LOW | (Rawls 2003; Leung et<br>al. 2016) |
| 1276227 | C | T | glycine receptor subunit<br>alpha-2 / gamma-<br>aminobutyric acid<br>receptor subunit alpha-6 | PF02932 \| PF2931 | 3 ' UTR | MODIFIER | - |
| 1404415 | G | C | omega-amidase NIT2 | PF00795 | 3 ' UTR | MODIFIER | - |
| 11174785 | A | G | 5-oxoprolinase | PF02538 \| PF05378 \|<br>PF01968 | 5 ' UTR | MODIFIER | - |
| 11690229 | A | T | glycine receptor subunit<br>alpha-2 | PF02931 | synonymous<br>variant | LOW | - |

## Discussion

### Genomic and transcriptomic variation data belong to a single *O. similis* lineage

Because genomes of small animals like copepods are difficult to reconstruct, we used *DiscoSNP++*, a reference-free variant caller to extract variants from metagenomic data, that already showed its good accuracy on *Tara* Oceans metagenomic data (Arif et al. 2018).

Global populations of *O. similis* are known to be composed of cryptic lineages across oceanic basins (Cornils et al. 2017). It is also known that this species in highly abundant among other copepods in Arctic Seas (Blachowiak-Samolyk et al. 2008; Dvoretsky 2012; Zamora-Terol et al. 2013; Castellani 2016). Thus, the assessment that the extracted variants from the seven samples used in our study belong to the same *O. similis* cryptic lineage was a prerequisite for further analyses. Three different analyses support this assumption. First, the distribution of depth of coverage in each of the seven samples followed the expected negative binomial distribution (Supplementary Figure S2). Indeed, the possibility to observe these patterns in the presence of different species would require them to be equally co-abundant, which is unlikely. Thus, this covariation of the depth of coverage of these variants supports the single species genome origin. Secondly, the high proportion of variants (97%) mapped on the Mediterranean *O. similis* transcriptomes, another cryptic lineage (Cornils et al. 2017), showed that the variant clustering method was efficient to regroup loci of *O. similis*. Finally, the unimodal distribution of *F*_ST_ showed that these populations of *O. similis* belong to the same polar cryptic species, and that all loci are under neutral evolution. Altogether, these results show that we were able to retrieve polymorphic data of a single species, *O. similis*, on which population genomics and psASE detection can be undertaken.

### *O. similis* populations are weakly structured within the Arctic Seas

We observed that the seven populations examined showed low genomic differentiation, despite the large distances separating them, which was illustrated by a non-significant Mantel test for isolation-by-distance (Figure 2a). *F*_ST_ and *pcadapt* analyses both showed the same patterns of genomic differentiation. Going into details, three different cases can be described. First, the differentiation of populations from TARA_155 and 178 is relatively high compared to the others. Secondly, the geographically close populations from TARA_210 and 209 present a relatively high differentiation (median pairwise-*F*_ST_ of 0.11, PC3). This could be explained by the West Greenland current acting as a physical barrier between the populations, which could lead to reduced gene flow (Myers et al. 2008). At last, the strong link between TARA_158 from Northern Atlantic current and TARA_206/208 from the Baffin Bay is the most intriguing. Despite the large distances that separate the first one from the others, these three populations are well connected. Based on this weak structure and that most of loci follows a neutral evolution (Figure 2d), outliers detected by *pcadapt* probably are truly under selection and not due to specific population differentiation.

Metagenomic data enable to draw the silhouette of the population genetics but lacks resolution when dealing with intra-population structure. However, our findings are concordant with previous studies underpinning the large-scale dispersal, interconnectivity of marine zooplankton populations in other oceans, at diverse degrees (Goetze 2005; Peijnenburg & Goetze 2013; Blanco-Bercial & Bucklin 2016; Höring et al. 2017). Weak genetic structure in the polar region was highlighted for other major Arctic copepods like *Calanus glacialis* (Weydmann et al. 2016), and *Pseudocalanus* species (Aarbakke et al. 2014). The absence of structure was explained by ancient diminutions of effective population size due to past glaciations (Bucklin & Wiebe 1998; Edmands 2001; Aarbakke et al. 2014), or high dispersal and connectivity between the present-day populations due to marine currents (Weydmann et al. 2016). Using Lagrangian time travel or dispersal probabilities could help to estimate how much marine currents explain this observed genomic differentiation.

### Population-scale ASE in *O. similis* populations

Usually, ASE analyses are achieved by measuring the difference in RNA-seq read counts of a heterozygous site at cellular, tissue or individual scales. Obtaining a comprehensive and exhaustive view of ASE at all these scales is merely possible due to sampling challenges for small organisms. Particularly, gathering a large number of individuals, sampling the tissues or organs of interest remain a technical barrier especially for uncultured small animals, when the amount of DNA retrieved from a single individual is not sufficient for high-throughput sequencing. Here, the detection of psASE was possible by comparing the observed allele frequencies based on metagenomic and metatranscriptomic data, which bypasses the obstacles previously described. The null hypothesis, supposing a strong correlation between the expression of an allele and its genomic abundance at the population-scale was confirmed in the seven populations, allowing us to continue and detect outliers targeted by psASE with our approach. In the case of psASE, we expect to detect strong patterns of unbalanced allelic expression present in a large fraction of the individuals of a population. Thus, a variant can be detected under psASE because of intra- or inter-individual ASEs. Intra-individual ASE detected at population-level should be present in the majority of individuals at cell or organ levels which constitutes the classical type of ASE mechanism. Inter-individual ASEs encompass ASEs due to physiological states, developmental stages, or sex differentiation. In *O. similis*, the proportion of each developmental stage and sex is known to vary within and between populations and across seasons (Lischka & Hagen 2005; Dvoretsky 2012).

In our study, the amount of detected psASE in each population was always lower than 1% of tested heterozygous variants, which altogether correspond to 2% of the total set of variants. In humans (Zhang et al. 2014), baboons (Tung et al. 2015) and flycatchers (Wang et al. 2017), 17%, 23% of genes and 7.5% of transcripts were affected by ASE respectively, meaning that psASE possibly tends to dilute individual ASE.

### Genomic drivers of population-scale ASE

The role of genomic background on psASE was deciphered by explaining the amount of shared psASEs by the genomic differentiation. From the strong correlation and the estimates of the model, we conclude that highly connected populations tend to share more loci under psASE than less connected populations, characterized by weak differentiation. However, even if only a small proportion of variants were targeted by psASE in different populations, our results show that genomic background partially drives psASE. To have a better understanding of the factors explaining psASE, environmental or population dynamics parameters (sex-ratio bias, proportions of larvae for example) could also be used as explanatory variables.

### Toward the link between population-scale ASE and natural selection

As genomic background among the seven populations partially explains psASE, we estimated the number of loci under psASE that were also targeted by natural selection. A significant amount of SNVs (152) were subject to selection among the seven populations and to psASE in at least one population, meaning that the two mechanisms can target the same genomic regions more than expected randomly.

Three main features of ASE can be under selection. First, the observed variant can be in linkage with another variation in upstream *cis*-regulatory elements like transcription factors fixation sites, or epialleles (Castel et al. 2015). Secondly, our annotation of candidate variants with SNPeff revealed a majority of variants located in 5’ and 3’UTRs, which are variations known to both affect transcription efficiency through mRNA secondary structures, stability and location (Mignone et al. 2002; Matoulkova et al. 2012; Dvir et al. 2013). For variants located in exons, a majority were identified as synonymous mutations, growingly described as potential target of selection by codon usage bias, codon context, mRNA secondary structure or transcription and translation dynamics (Ingvarsson 2010; Shabalina et al. 2013). Also, fifteen missense mutations were spotted, but with moderate predicted impact on protein amino acid composition. Nevertheless, we did not find premature nonsense mutation, even if variants under ASE has been described to trigger or escape potential nonsense-mediated decay, but the possibility that the causal variation is located in introns cannot be ruled out. However, due to the large possible origins of ASE and the lack of a proper reconstructed *O. similis* genome and genotypes, no major cause was identified from this study.

The process of adaptation through gene expression was studied in human populations and investigated thanks to the large amount of accessible data. In a first study, a link has been established between gene expression and selection, affecting particular genes and phenotypes, looking at *cis*-acting SNPs (Fraser 2013). In a second study, the team was able to detect loci under ASE and selection at the same time in different human populations (Tian et al. 2018). Finally, approaches in a plant model, *Capsella grandiflora*, a species characterized by weak population structure and large effective population size, emphasized the relative impact of purifying selection and positive selection on *cis*-regulatory variation in populations (Josephs et al. 2015; Steige et al. 2017). In our results, we identified a large fraction of candidate loci (Supplementary Figure S5) being under psASE in one population with a very low genomic frequency and tending to fixation in other populations (as exampled in Figure 4c). This could be linked to an adaptation of a population to environmental changing conditions during constant oceanic migration, leading to a purifying selection on the considered allele or a positive selection on the alternative allele. Our observations thus complete previous analyses, as they quantify the link between psASE and selection in natural marine populations composed of many individuals and reveal the evolutive potency of ASE, for the first time at the population-scale. Together, these results show that the two mechanisms lead to a similar molecular phenotype in allelic expression. This raises questions on whether ASE and selection are independent or sequential mechanisms.

### Nervous system and visual perception are important targets of the natural selection and population-scale ASE in *Oithona similis*

This evolutive link between psASE and selection is supported by the biological functions associated to the targeted genes, which are involved notably in the copepods nervous system in two ways. The first result is the presence of genes implicated in glutamate metabolism and glycine and/or GABA receptors. Glutamate and GABA are respectively excitatory and inhibitory neurotransmitters in arthropods motor neurons (Smarandache-Wellmann 2016). Glycine and GABA receptors have already been described as a target of selection in *Oithona nana* in Mediterranean Sea (Madoui et al. 2017; Arif et al. 2018). Secondly, the functional analysis revealed also the importance of the eye and visual perception in the *O. similis* evolution.

Together, this shows that psASE has an evolutive impact on functions linked to the nervous system of copepods. The latter constitutes a key trait for its reproduction and survival, and based on our data and previous work, a prime target for evolution, allowing higher capacity of perceiving and fast reacting leading to more efficient predator escape, prey catching and mating. This can explain the great evolutive success of these animals (Svensen 2000; Kiørboe et al. 2010; Kiørboe 2011).

## Conclusion

Gene expression variation is thought to play a crucial role in evolutive and adaptive history of natural populations. Herein, we integrated metagenomic and metatranscriptomic data to detect ASE at the population level for the first time. Then, we demonstrated the link between psASE and genomic background of the populations on one hand and with natural selection on the other hand, by providing a quantitative observation of this phenomenon and its impact on specific biological features of copepods. In the future, we will try to expand these observations to other organisms and question the nature of the link between psASE and natural selection.

## Acknowledgments

This work was supported by sponsors who participated in the *Tara* Oceans Expedition 2009–2013: Centre National de la Recherche Scientifique, European Molecular Biology Laboratory, Genoscope/Commissariat à l’Energie Atomique, the French Government “Investissements d’Avenir” programmes OCEANOMICS (ANR-11-BTBR-0008), FRANCE GENOMIQUE (ANR-10-INBS-09-08), Agnes b., the Veolia Environment Foundation, Region Bretagne, World Courier, Illumina, Cap L’Orient, the Electricite de France (EDF) Foundation EDF Diversiterre, Fondation pour la Recherche sur la Biodiversite, the Prince Albert II de Monaco Foundation, Etienne Bourgois and the *Tara* schooner and its captain and crew. *Tara* Oceans would not exist without continuous support from 23 institutes (oceans.tara-expeditions.org). We acknowledge the support of Vincent Segura and Leila Tirichine for fruitful scientific discussions and support on the analyses and manuscript. This is contribution number XX from *Tara* Oceans.

## Author’s contributions

Individuals for transcriptome production were sampled by J-LJ and KS. KS extracted RNA, EP and KL prepared the libraries and sequencing, MAM assembled the reads and RLJ annotated transcriptomes. PP and CA gave expertise support on *DiscoSNP++* and statistical framework respectively. RLJ and MAM performed the analyses and wrote the manuscript. MAM designed and supervised the study. J-LJ and PW offered scientific support.

## Tables

**Supplementary Table S1:** *Oithona similis* Mediterranean transcriptomes summary

**Supplementary Table S2:** *Tara* Oceans and *Oithona similis* Mediterranean transcriptomes samples accession numbers

**Supplementary Table S3:** Variants dataset

**Supplementary Table S4:** Variants targeted by ASE and selection functional annotations

**Supplementary Table S5:** Variant annotation by SNPeff

## Figures

**Supplementary Fig S1:** Validation of taxonomic assignation

**Supplementary Fig S2:** *O. similis* depth of coverage of biallelic loci in seven *Tara* Oceans samples

**Supplementary Fig S3:** Population genomic differentiation

**Supplementary Fig S4:** Genomic differentiation and geographic distance

**Supplementary Fig S5:** Metagenomic and metatranscriptomic profiles of candidate loci

**Supplementary Fig S6:** Functional analysis of *O. similis* transcripts targeted by ASE and selection

