## Supplementary for "Investigating Population-scale Allele Specific Expression in Wild Populations of *Oithona similis* (Cyclopoida, Claus 1866)"

### Supplementary Figures

**Supplementary Figure 1 : Validation of taxonomic assignation.** In rows are represented the 82 accession numbers of *Oithona* species 28S sequences. In bold, type localities of *O. similis* as described in Cornils et al., 2017. In columns are represented ribosomal read sets of the eight individuals.

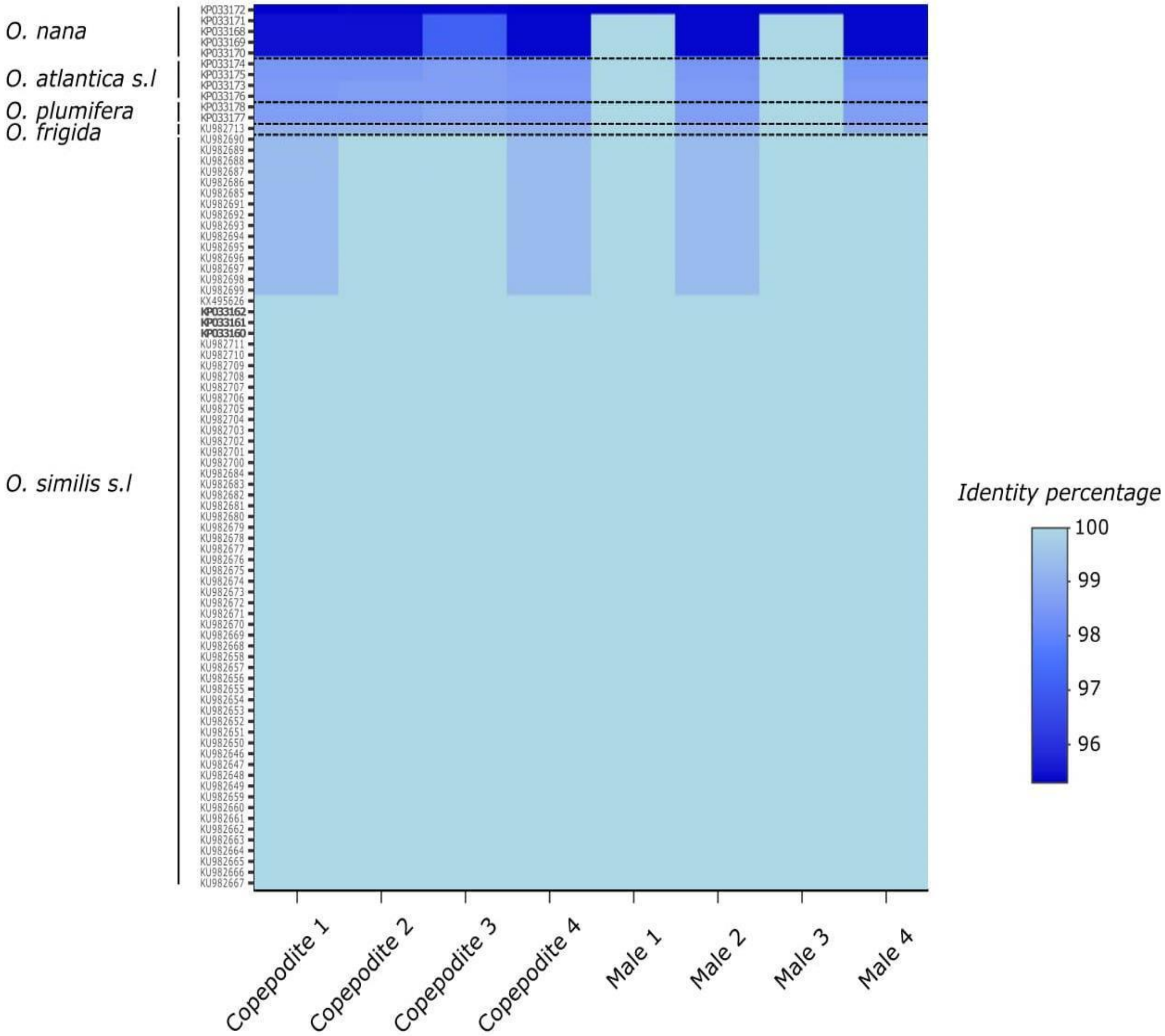

**Supplementary Figure 2 : *Oithona similis* depth of coverage of biallelic loci in seven Tara Oceans samples.** In red is represented the theoretical depth of coverage. In black is represented the observed depth of coverage. **a,** Comparison of coverage distribution. **b,** Comparison of the cumulative distribution functions.

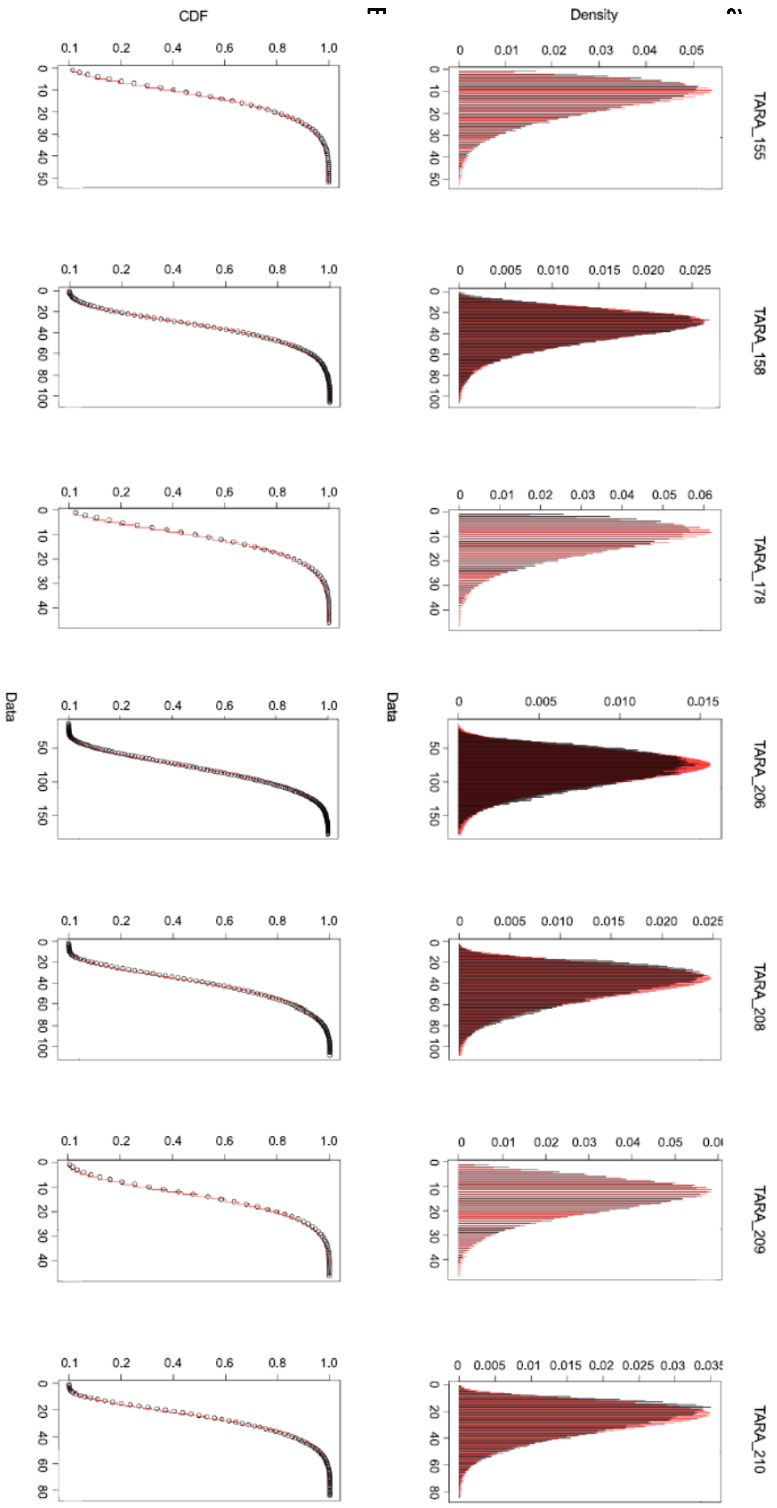

**Supplementary Figure 3 : Population genomic differentiation. a to c,** Principal Component Analysis from *pcadapt*. Each axis is a principal component with the corresponding proportion of variance explained between brackets **d,** Pairwise-$F_{ST}$ distributions.

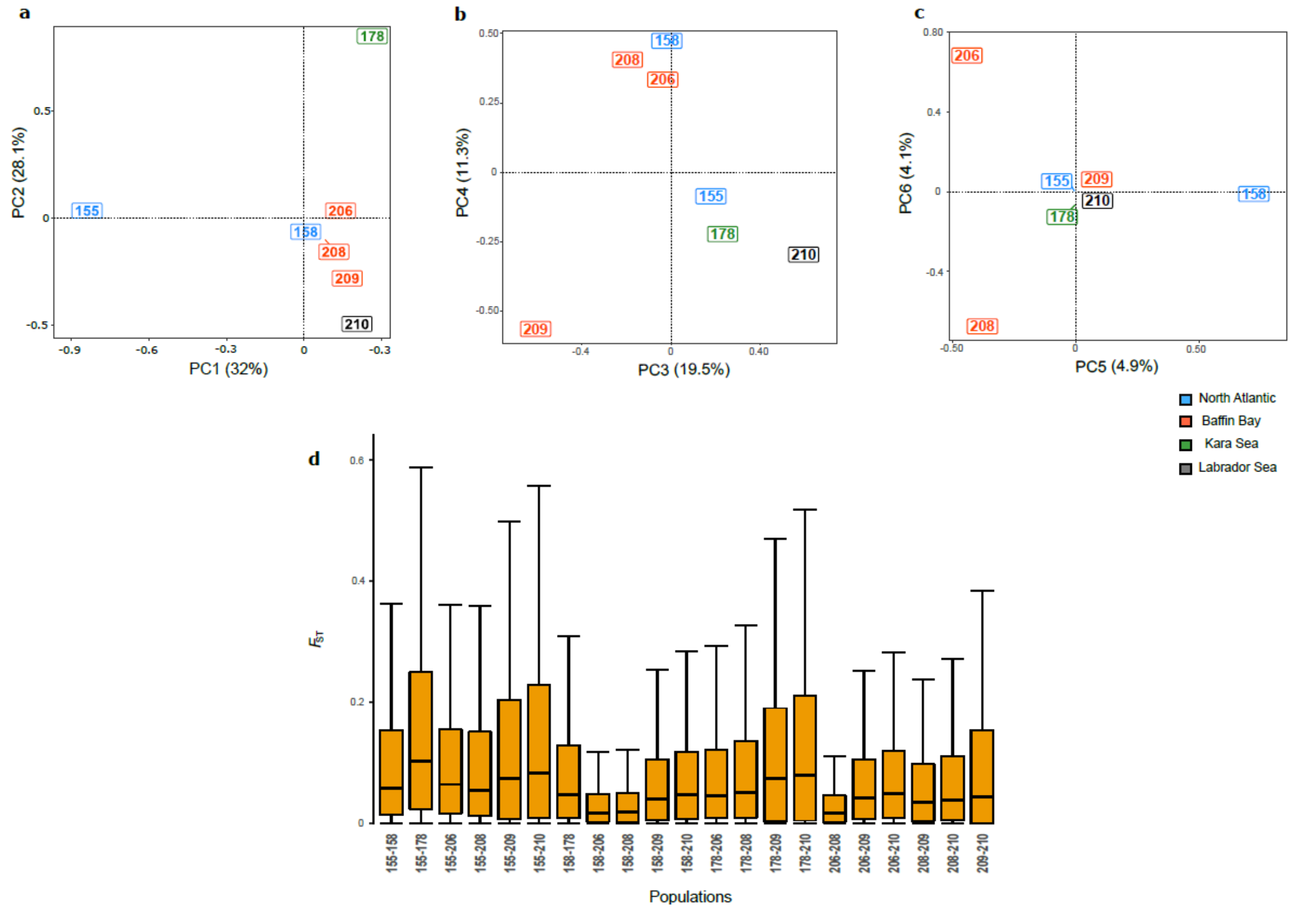

**Supplementary Figure 4 : Genomic differentiation and geographic distance.**
Plot displaying Pairwise-$F_{ST}$ and corresponding geographic distance. In blue, linear regression curve. In grey, the 95% confidence interval.

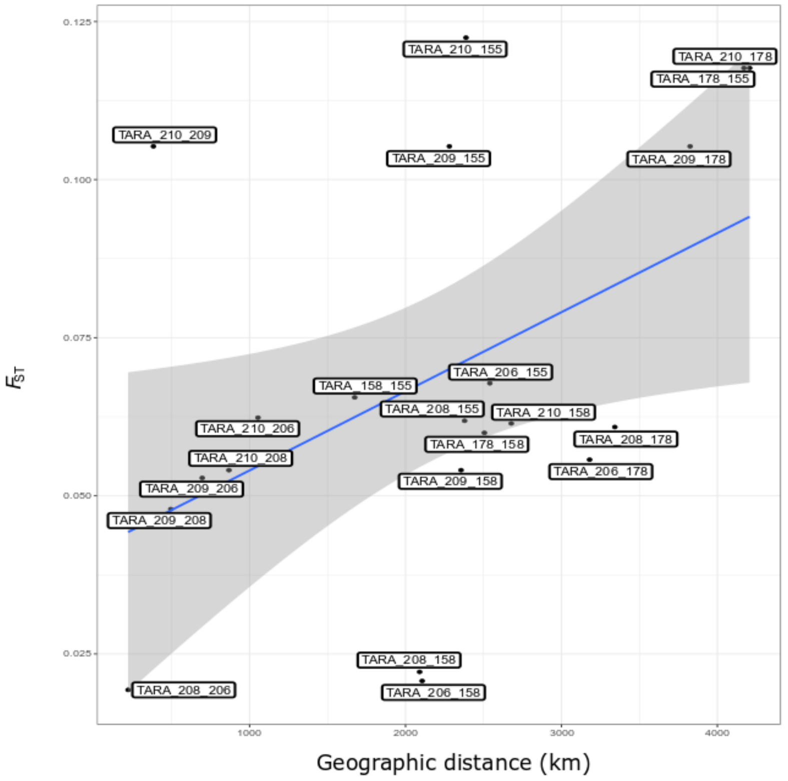

**Supplementary Figure 5 : Metagenomic and metatranscriptomic profiles of candidate loci.** Each graph represents BAF (gold) and BARE (blue) in the seven population of a candidate variant under psASE and selection. Asterisks above the barplot means the variant has been detected under psASE in the corresponding population (Deviation test corrected p-value < 0.1*, < 0.05**, <0.01***, <0.001****).

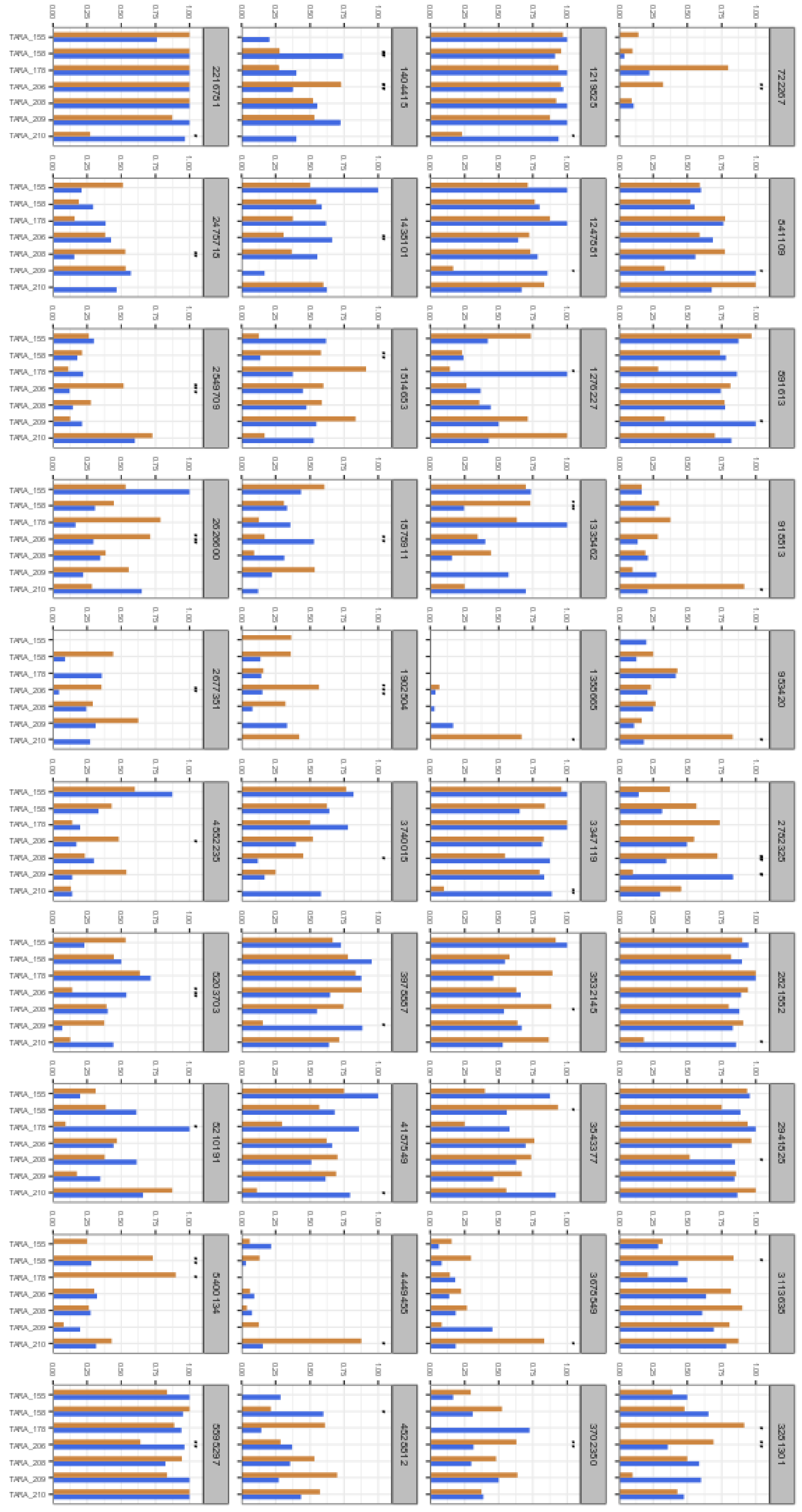

8

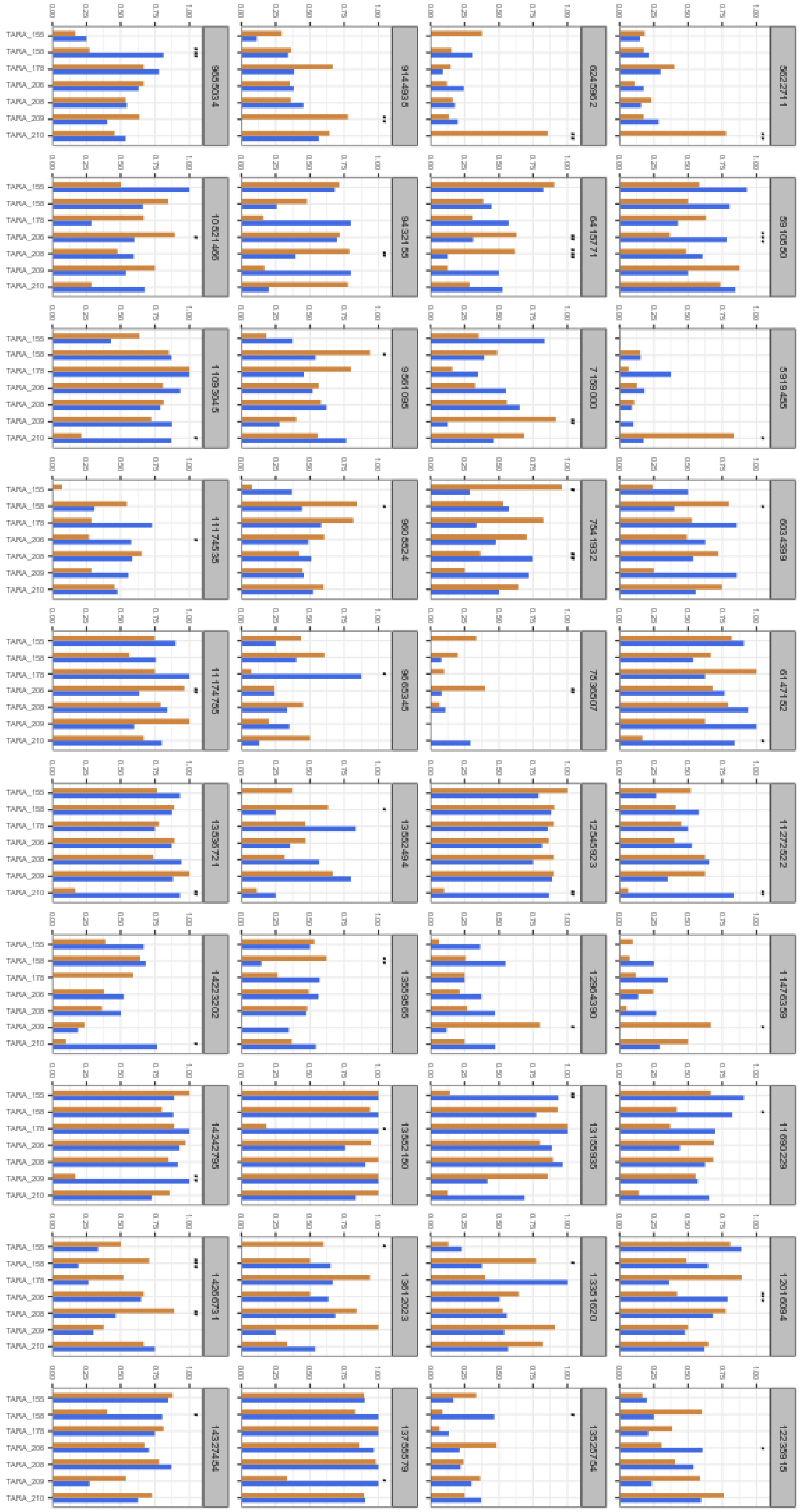

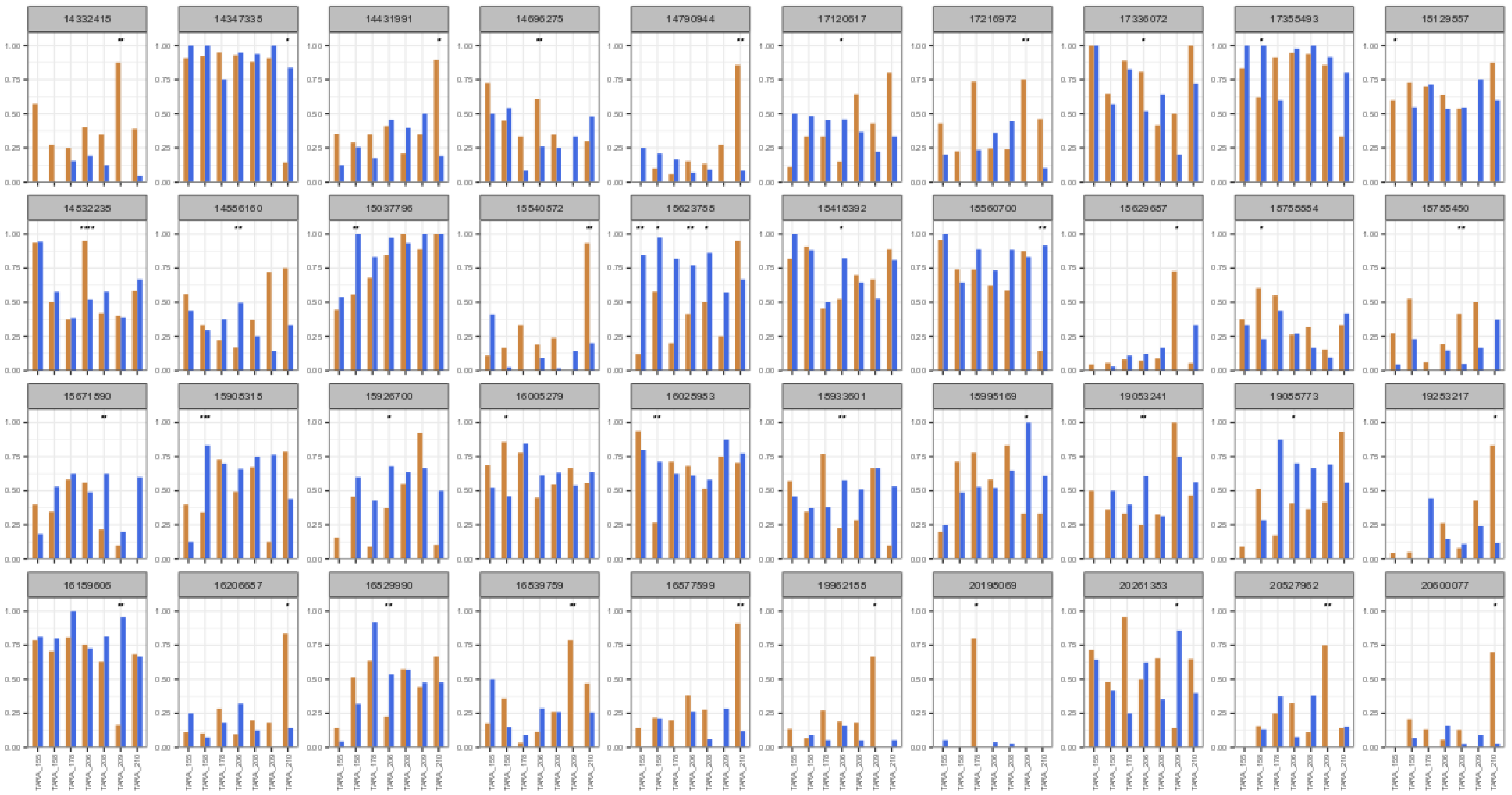
14332415
14347335
14431991
14696275
14790944
17120617
17216972
17336072
17355493
18129857
14532235
14556160
15037796
15540572
15623755
15415392
15560700
15629657
15755554
15755450
15671590
15905315
15926700
16005279
16025953
15933601
15995169
19053241
19055773
19253217
16159606
16206657
16529990
16539759
16577599
19962155
20195069
20261353
20527962
20600077
0.00
0.25
0.50
0.75
1.00
TARA_155
TARA_158
TARA_178
TARA_206
TARA_208
TARA_209
TARA_210

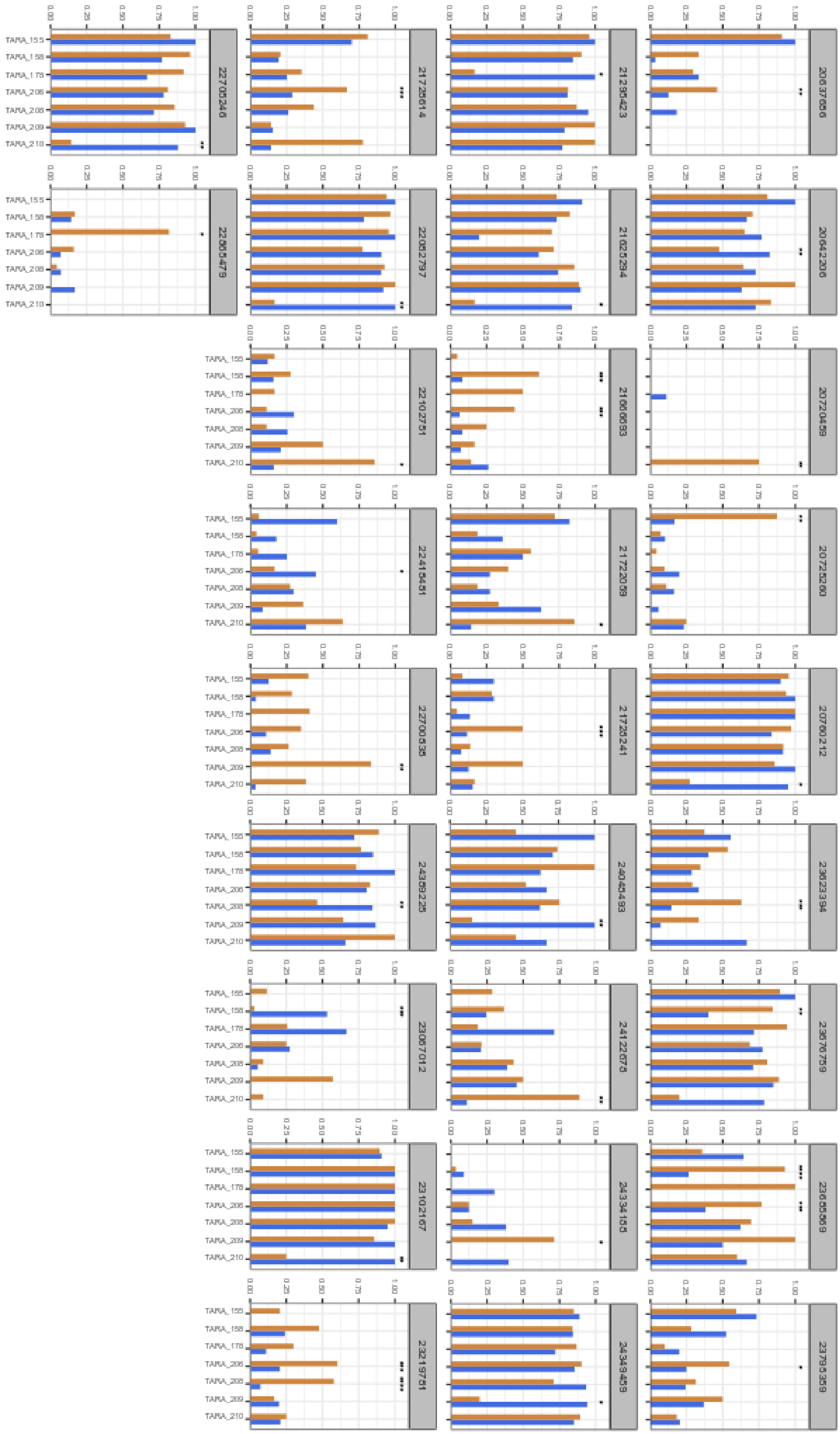
0.00
0.25
0.50
0.75
1.00
TARA_155
TARA_158
TARA_178
TARA_206
TARA_208
TARA_209
TARA_210
22705246
22865479
21728614
22082797
22102751
22415451
22700838
24358225
23067012
23102167
23219751
21235423
21625294
21666083
21722058
21726241
24045483
24122678
24334155
24349459
20637656
20642206
20720459
20725260
20760212
23623394
23676759
23685589
23795359

**Supplementary Figure 6 : Functional analysis of *Oithona similis* transcripts targeted by ASE and selection.** The axes correspond to statistical metrics computed by *dcGO Enrichment*. Are highlighted the most significantly enriched terms in **a,** Biological Process GO-terms, **b,** Cellular Component GO-terms, **c,** Molecular Functions GO-terms.

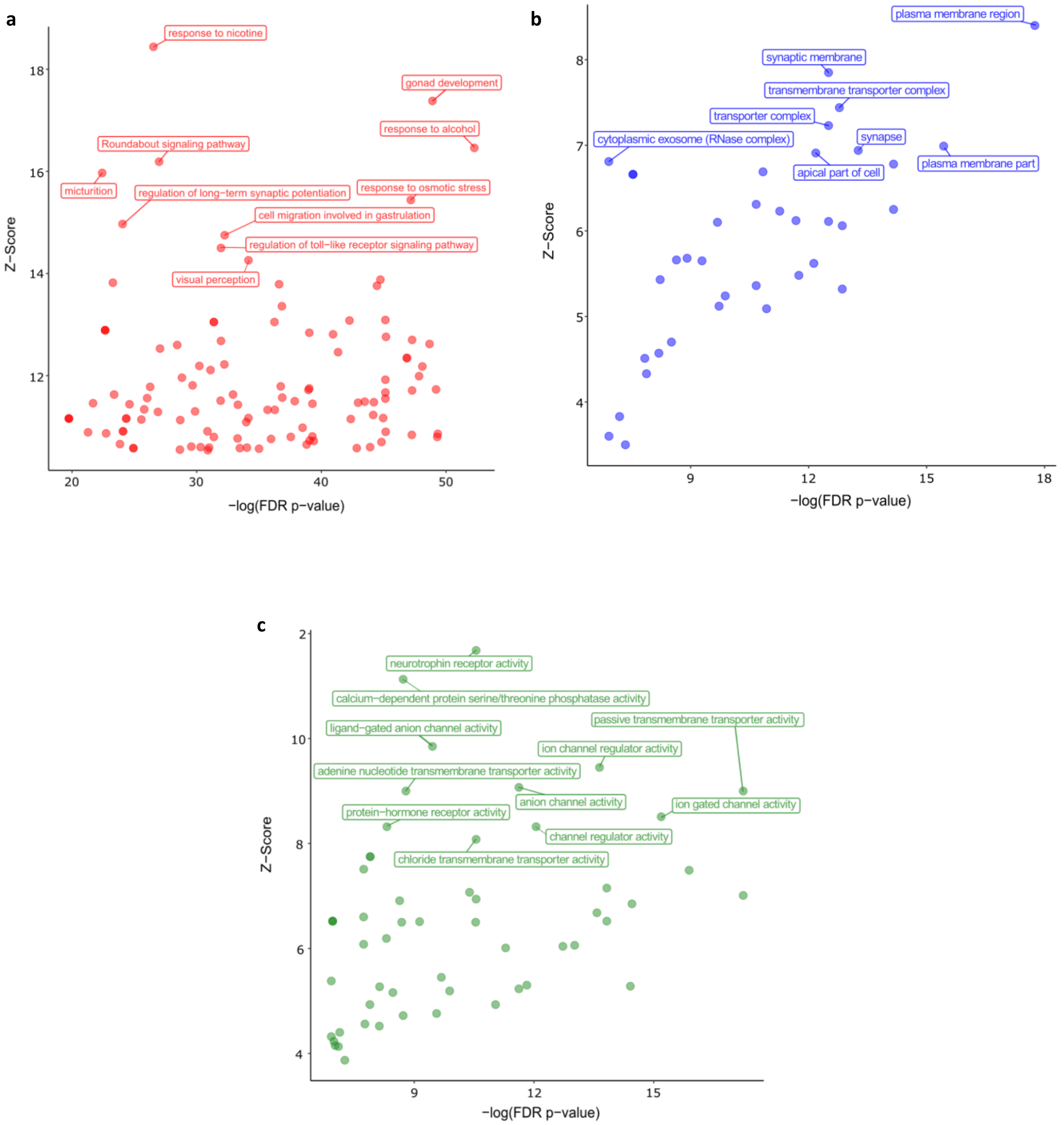

**Supplementary Table S1 : *Oithona similis* Mediterranean transcriptomes summary.**

| | Copepodite | | | | Male | | | |
|---|---|---|---|---|---|---|---|---|
| | **1** | **2** | **3** | **4** | **1** | **2** | **3** | **4** |
| **Number of reads** | 18,052,409 | 21,048,656 | 19,129,821 | 17,624,872 | 17,823,526 | 19,237,768 | 18,292,442 | 13,933,841 |
| **Number of transcripts** | 81,802 | 82,811 | 131,747 | 83,244 | 117,951 | 79,938 | 115,170 | 79,038 |
| **Number of transcripts with ORF prediction** | 20,389 | 20,963 | 42,404 | 20,746 | 38,580 | 19,488 | 34,179 | 20,482 |

**Supplementary Table S5 : Variant annotation by SNPeff**. In bold and with an asterisk, significant values under hypergeometric test.

| SNPeff prediction | Variants under ASE | Variants under selection | Variants under ASE & selection | Total |
|---|---|---|---|---|
| Missense | **54* (12.4%)** | **66* (12.6%)** | 15 (9.9%) | 2529 (9.8%) |
| Synonymous | 170 (39.1%) | 213 (40.8%) | 59 (38.8%) | 11999 (46.6%) |
| Start lost | 0 | 0 | 0 | 4 (0.02%) |
| Stop gained | 1 (0.2%) | 0 | 0 | 4 (0.02%) |
| Stop retained | 0 | 0 | 0 | 2 (0.01%) |
| 3'UTR | 75 (17.2%) | **111* (21.3%)** | 31 (20.4%) | 4480 (17.4%) |
| 5'UTR | 80 (18.4%) | 79 (15.1%) | 29 (19.1%) | 4110 (16%) |
| 5'UTR premature start | 12 (2.8%) | 16 (3.1%) | 3 (2%) | 778 (3%) |
| No Transdecoder Hit | 23 (5.3%) | 20 (3.8%) | 8 (5.3%) | 1142 (4.4%) |
| Not mapped | 20 (4.6%) | 17 (3.3%) | 7 (4.6%) | 720 (2.8%) |
| TOTAL | 435 | 522 | 152 | 25768 |
